# Polycystin-1 C-Terminus Regulates Protein Synthesis-Related Pathways in Cardiomyocytes

**DOI:** 10.64898/2026.03.21.713243

**Authors:** Matthew Fiedler, Alejandra Vasquez Limeta, Ernesto Reyes-Sanchez, Monserrat Reyes-Lozano, William Perez, Leah Carter, Christopher J. Ward, Francisco Altamirano

## Abstract

Pathologic cardiac hypertrophy requires increased protein synthesis, but the mechanosensors that link membrane stretch to translational control remain poorly understood. Polycystin-1 (PC1), encoded by *PKD1*, has been proposed as a cardiac mechanosensor, with its C-terminal tail (PC1-CT) promoting hypertrophy in rodent cardiomyocytes. However, its subcellular localization and downstream signaling remain incompletely defined, especially in human cardiomyocytes.

Here, we examined endogenous PC1 C-terminus localization and the effects of adenoviral PC1-CT overexpression in human iPSC-derived ventricular cardiomyocytes (hiPSC-CMs) and adult mouse ventricular myocytes. Immunofluorescence revealed a striking striated pattern for both endogenous PC1 C-terminus (detected with a PC1-CT antibody) and the overexpressed PC1-CT fragment. In hiPSC-CMs, the PC1 C-terminus localized between the α-actinin bands. In contrast, in adult cardiomyocytes, the overexpressed protein colocalized with α-actinin and desmin, suggesting that PC1-CT sarcomeric distribution depends on cardiomyocyte maturation.

We performed RNA-seq to assess transcriptional responses downstream of PC1-CT overexpression in hiPSC-CMs relative to LacZ controls. Gene Set Enrichment Analysis (GSEA) revealed enrichment of gene sets related to ribosome biogenesis, RNA processing, and protein synthesis, while classical hypertrophic markers remained unchanged. Pathway analysis suggested increased PI3K activity. PC1-CT overexpression increased phosphorylation of Akt, ERK, S6K1, and ribosomal protein S6 without altering 4EBP1 phosphorylation, suggesting preferential activation of the mTOR-S6K1-S6 branch. Pharmacological studies showed that pan-PI3K inhibition abolished S6 phosphorylation, whereas MEK blockade did not affect it; pertussis toxin and PI3Kγ-selective inhibitors also did not affect S6, suggesting a G_i/o_-independent PI3K/Akt signaling driving mTOR-S6K1-S6 activation.

Collectively, these data identify a sarcomere-associated pool of PC1-CT that engages PI3K-Akt-mTOR-S6K1-S6 signaling to enhance transcriptional programs related to ribosome biogenesis and protein synthesis, without activating a canonical hypertrophic gene program. These findings reveal a mechanistic link between PC1-CT and cardiomyocyte growth.

## INTRODUCTION

Pathologic cardiac hypertrophy is a maladaptive response to mechanical overload and a major determinant of progression to heart failure [1]. Cardiomyocyte growth depends on increased protein synthesis, but the molecular sensors that couple membrane stretch to this process remain poorly defined [2]. Polycystin-1 (PC1), a large transmembrane protein encoded by *PKD1*, has been implicated in cardiac hypertrophy and proposed to act as a mechanosensor in cardiomyocytes [3]. In rodent models, loss of PC1 blunted both *in vitro* stretch-induced cardiomyocyte hypertrophy and *in vivo* pressure overload-induced hypertrophy [3].

Loss of PC1 resulted in proteasomal degradation of the L-type Ca^2+^ channel (LTCC) following *in vitro* mechanical stretch. In contrast, overexpression of the soluble cytoplasmic C-terminal tail of PC1 (PC1-CT) stabilized LTCC, enhanced protein synthesis, and increased cardiomyocyte hypertrophic markers [3]. A follow-up study showed that a synthetic membrane-anchored PC1-CT construct – containing the extracellular domain and signal peptide of CD16 and the transmembrane domain of CD7 – enhanced Akt activation via its G-protein domain, thereby increasing LTCC content [4]. These studies indicate that PC1-CT is an important mediator of the cardiomyocyte response to mechanical stress.

Cardiac hypertrophy is a complex response, and the mechanisms by which PC1-CT regulates protein synthesis and hypertrophy remain largely unknown. Moreover, PC1-CT is known to regulate signaling at the plasma membrane through a heterotrimeric G-protein activation sequence [5], or after cleavage via nuclear and mitochondrial localization sequences [6, 7], a coiled-coil domain [8], and other putative interaction domains. However, PC1-CT subcellular distribution, downstream signaling, and effects on gene expression in cardiomyocytes remain unaddressed [9].

Here, we show that endogenous PC1 C-terminus and overexpressed PC1-CT display a striated, predominantly sarcomeric distribution in hiPSC-CMs and adult mouse ventricular cardiomyocytes, with limited overlap with intracellular sarco-endoplasmic reticulum KDEL marker. Transcriptional profiling after adenoviral PC1-CT overexpression identified enrichment of ribosome biogenesis and protein synthesis-related gene pathways, together with activation of mTOR downstream effectors, without changes in classical hypertrophic gene markers.

## METHODS

### hiPSC differentiation

hiPSC line hPSCreg #TMOi001-A (Gibco #A18945) was grown in mTESR Plus (STEMCELL Technologies #100-0274 and #100-0275), and differentiated into cardiomyocytes, selected, and then expanded [10]. Cells were used for experimentation after removal of CHIR-99021 and a two-week recovery period in culture. Experiments were conducted after day 35, when the ventricular cardiomyocyte population is known to increase. For localization experiments, triiodothyronine and dexamethasone were used to promote further maturation [11].

### Adult cardiomyocyte isolation

All procedures were approved by the Houston Methodist Research Institute IACUC and were conducted in accordance with NIH guidelines for animal care. Animals were anesthetized with 5% isoflurane and cervically dislocated under deep anesthesia. Hearts were quickly dissected and retrogradely perfused with Liberase TM and trypsin, then cells were filtered, and Ca^2+^ was gradually reintroduced as described [12].

### Adenovirus infections

Adenoviruses expressing LacZ or PC1-CT fused to a V5 tag (amino acids 4102-4303) have been described previously [12]. The Adeno-X^™^ Adenoviral System 3 (Takara) was used to generate adenoviruses expressing mCherry or mCherry-PC1-CT fused to a V5 tag. hiPSC-CMs and adult cardiomyocytes were infected with a multiplicity of infection (MOI) of 10 and lysed at 48 h for RNA-seq and 72 h for staining.

### Immunofluorescence

hiPSC-CMs were fixed using 4% paraformaldehyde and permeabilized with 0.1% Triton X-100 in phosphate-buffered saline (PBS). Cells were blocked with 5% normal goat serum (NGS) and 3% bovine serum albumin (BSA) in PBS. Isolated adult mouse ventricular cardiomyocytes were fixed using 100% methanol at -20°C. Permeabilization and blocking were performed simultaneously with 5% NGS in 0.5% Triton X-100 in PBS. Primary antibodies mouse anti-human PC1-CT (161F, 1:200) [13], rabbit anti-α-actinin-2 (1:200, Invitrogen, #701914), rabbit anti-mCherry antibody (1:200, Cell Signaling, #43590), mouse anti-α-actinin2 (1:200, Invitrogen, #MAI-22863), mouse anti-desmin (1:100, Invitrogen, #MAS-13259), and rabbit anti-KDEL (1:100, Invitrogen, #PA1-013). All primary antibodies were incubated overnight at 4 °C in blocking buffer. After washing, fluorophore-conjugated secondary antibodies (goat anti-mouse or anti-rabbit Alexa Fluor 488 or 555) were used, followed by DAPI staining. Images were collected using a Nikon AX NSPARC confocal microscope. Python scripts were used for figure generation, ROI selection, and x-y intensity profile analysis. For visualization, fluorescence intensities were rescaled using percentile-based min-max normalization (2^nd^-98^th^ percentile) and smoothed with a Gaussian filter (σ = 0.3 px). When indicated, selected zoomed ROIs were renormalized locally. Images are displayed as single-channel grayscale panels or pseudocolor overlays with scale bars. For intensity profile plots, raw intensities were sampled with a 5-pixel line width, smoothed along the profile (σ = 1 px), baseline-corrected with a rolling-minimum filter, and normalized to 0-1 using the 1^st^-99^th^ percentile, except in the siRNA comparison, where raw profiles were plotted directly.

### siRNA knockdown

hiPSC-CMs were transfected with control (SIC001, Universal Control, Mission Sigma) or PC1 (SASI_Hs01_00136684/PKD1) using Lipofectamine RNAiMAX (Thermo Fisher). Cells were stained after 72 h to determine 161F staining specificity.

### Western blots

Cells were lysed using 2X Laemmli buffer (Bio-Rad) supplemented with dithiothreitol (50 mM). Proteins were resolved using 4-20% TGX gels (Bio-Rad) and transferred to nitrocellulose membranes. Primary antibodies were incubated overnight at 4°C, and fluorophore-conjugated secondary antibodies were used for detection. The following primary antibodies were obtained from Cell Signaling and used at 1:1000: pERK (#4370), ERK (#9107), pAkt (#4060), Akt (#2920), pS6K1 (#9234), S6K1 (#2708), pS6 (#4858), S6 (#2317), p4EBP1 (#2855), non-phospho 4EBP1 (#4923) and total 4EBP1 (#9644). The following antibodies were obtained from Bio-Rad: V5 (#MCA1360, 1:2000) and hFAB™ Rhodamine anti-GAPDH (#12004167, 1:5000). Secondary antibodies IRDye 800CW (anti-rabbit, Li-COR) and StarBright 700 (anti-mouse, Bio-Rad) were used for detection using Chemidoc MF and analyzed with ImageLab software (Bio-Rad).

### RNA-seq

Total RNA was extracted using Quick-RNA Miniprep (Zymo). RNA quality, integrity, library preparation, next-generation sequencing, and FASTQ generation were performed by GeneWiz; all samples had an RNA integrity number> 9.4. Libraries were prepared using the NEBNext Ultra II RNA Library Prep Kit for Illumina (NEB). Poly(A) mRNA was enriched with Oligo(dT) beads, fragmented, and reverse-transcribed to cDNA. After end repair, A-tailing, and ligation of universal adapters, indexed libraries were amplified by limited-cycle PCR, validated on TapeStation, and quantified by Qubit and qPCR. Sequencing was performed on an Illumina HiSeq instrument with 2 × 150bp paired-end reads. Base calling used HiSeq Control Software (HCS); bcl files were converted to FASTQ and de-multiplexed with bcl2fastq 2.17. One mismatch was allowed for index sequence identification. FASTQ quality was assessed with FastQC (v0.12.1), trimmed with fastp (v0.24.0), and transcript counts obtained with Salmon (v1.10.0). Differential expression analysis was employed using RStudio 2025.9.2.418, R4.5.2, DESeq2 (1.50.2), and clusterProfiler (4.18.4) for Gene Set Enrichment Analysis (GSEA) [14] using GO Biological Processes gene sets. Pathway inference was done using decoupleR (2.16.0) [15].

## Data availability

Preliminary differential expression analysis outputs generated by GeneWiz were previously archived in Zenodo (DOI: 10.5281/zenodo.10901271). Because that repository does not include the complete RNA-seq dataset, the raw data and normalized counts underlying this study will be deposited in GEO before manuscript acceptance. The results presented here are based on reanalysis of FASTQ files using the workflow described above and are consistent with earlier findings.

### Statistics

Data are presented as box plots showing the median and interquartile range (IQR). Normality was assessed with the Shapiro-Wilk test. Because most datasets were not normally distributed, group comparisons were performed using the Mann-Whitney test. ANOVA was not used because our goal was to test specific pairwise comparisons within the same group. Statistical analyses were conducted in RStudio 2025.9.2.418 and with R 4.5.2.

## RESULTS

### Endogenous PC1 C-terminus and overexpressed PC1⍰CT exhibit a predominantly sarcomeric distribution in cardiomyocytes

To determine endogenous PC1 C-terminal localization, we immunolabeled hiPSC-CMs with a human-specific anti-PC1-CT antibody [13]. Confocal imaging revealed a consistent striated pattern resembling sarcomere organization. This signal was reduced by PC1 siRNA knockdown (**Fig. 1A**). Co-immunostaining with α-actinin showed a regular alternating pattern with a periodicity of ∼1.7 µm for both signals (**Fig. 1B**). To test whether this striated pattern overlapped with the sarco-endoplasmic reticulum, we used a KDEL antibody (**Fig. 1C**). The limited overlap between PC1 C-terminus and KDEL, together with the absence of mature T-tubules in hiPSC-CMs [16], supports a predominantly sarcomeric distribution. This localization was reproduced in hiPSC-CMs expressing mCherry-PC1-CT (PC1 amino acids 4102-4303) compared with mCherry controls; some cells additionally displayed costamere-like and nuclear localization depending on expression levels (**Fig 1D**). Live cell imaging of adult mouse ventricular cardiomyocytes infected with mCherry-PC1-CT showed a similar distribution (**Fig. 1E**). In adult cardiomyocytes, coimmunolabeling with α-actinin and desmin showed greater overlap of overexpressed PC1-CT with both markers (**Fig. 1F**).

**Figure 1.**
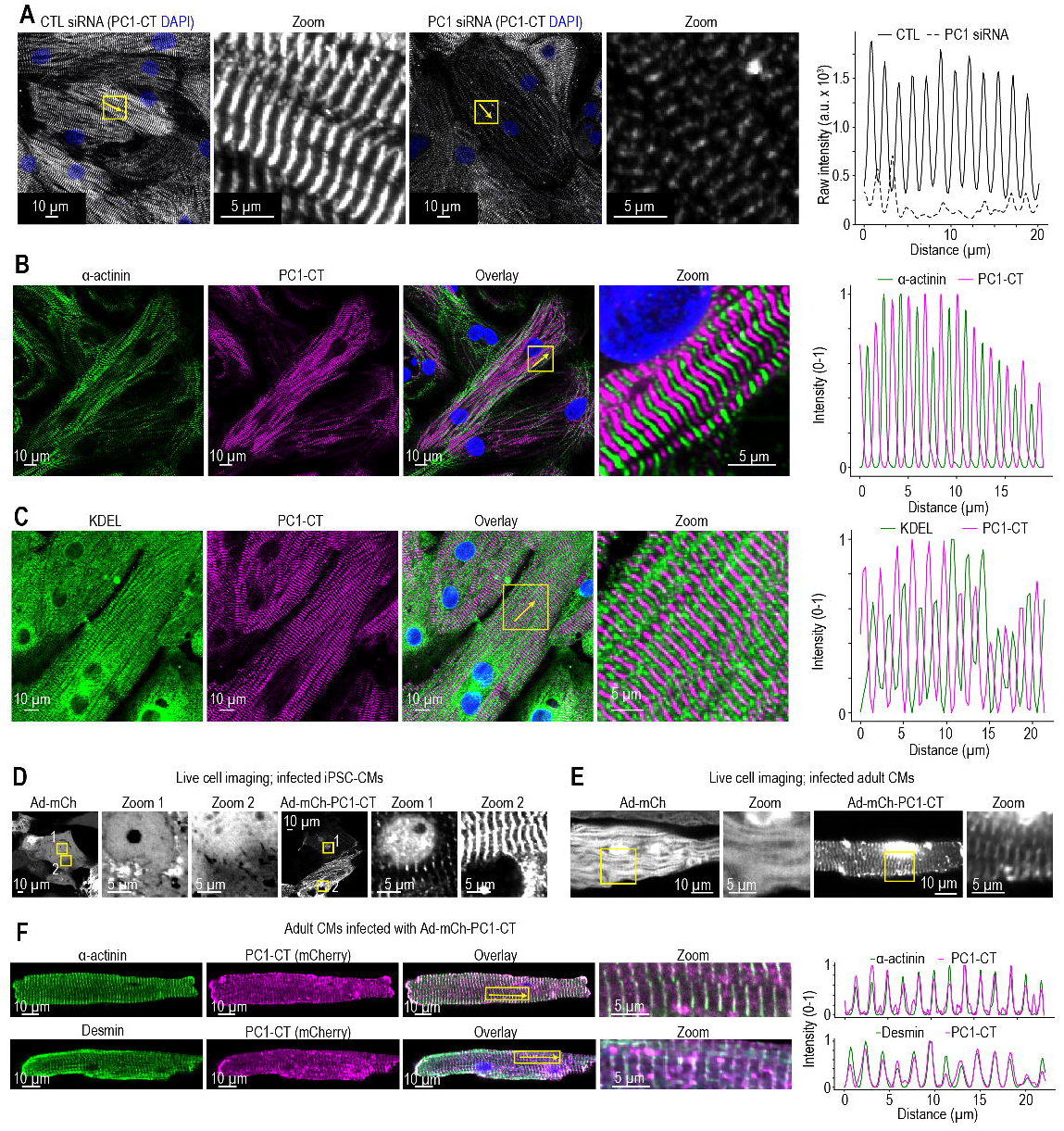
Localization of endogenous PC1 C-terminus and overexpressed PC1-CT. **A**. Maximum z-projected images from immunofluorescence confirm endogenous PC1 C-terminal localization with a striated pattern that resembles cardiomyocyte sarcomeres (white, 161F antibody) in hiPSC-CMs. siRNA-mediated PKD1 knockdown demonstrates antibody specificity. DAPI (blue) is used to label nuclei in all panels except panel **D-E**. Panels **B–F** are single confocal optical sections. The far-right panels show the raw fluorescence intensity profile for panel **A** and the normalized fluorescence intensity profiles for panels **B, C**, and **F**. **B**. Endogenous PC1 C-terminus (magenta) localizes in a striated pattern alternating with α-actinin staining (green), consistent with sarcomeric alignment. **C**. Endogenous PC1 C-terminus (magenta) shows minimal colocalization with the sarco-endoplasmic reticulum marker (KDEL, green). **D, E**. Representative live-cell images of hiPSC-CMs and adult mouse ventricular CMs expressing viral constructs encoding mCherry or mCherry-PC1-CT. PC1-CT localizes with a striated pattern and, in some cells, nuclear localization (zoom panels were renormalized to highlight localization). **F**. Localization of mCh-PC1-CT in adult vCMs aligns with Z-line proteins α-actinin (upper panel) and desmin (lower panel).

### PC1-CT regulates ribosome biogenesis and gene programs related to protein synthesis

RNA-seq after adenoviral PC1-CT overexpression in hiPSC-CMs identified genes that were differentially expressed relative to LacZ controls (48 h, MOI of 10; **Fig. 2A**). GSEA showed enrichment of ribosome biogenesis, RNA processing, nuclear export, and protein synthesis-related gene sets (**Fig. 2B**). Classical hypertrophic markers, however, were unchanged by PC1-CT overexpression (**Supplemental Table 1**). To infer upstream pathway activity, we applied PROGENy, implemented in decoupleR, using DESeq2 Wald statistics [15]. This analysis showed increased PI3K signaling in PC1-CT-expressing versus LacZ-expressing cardiomyocytes (**Fig. 2C**), supported by changes in downstream PI3K-responsive genes (**Fig. 2D**).

**Figure 2:**
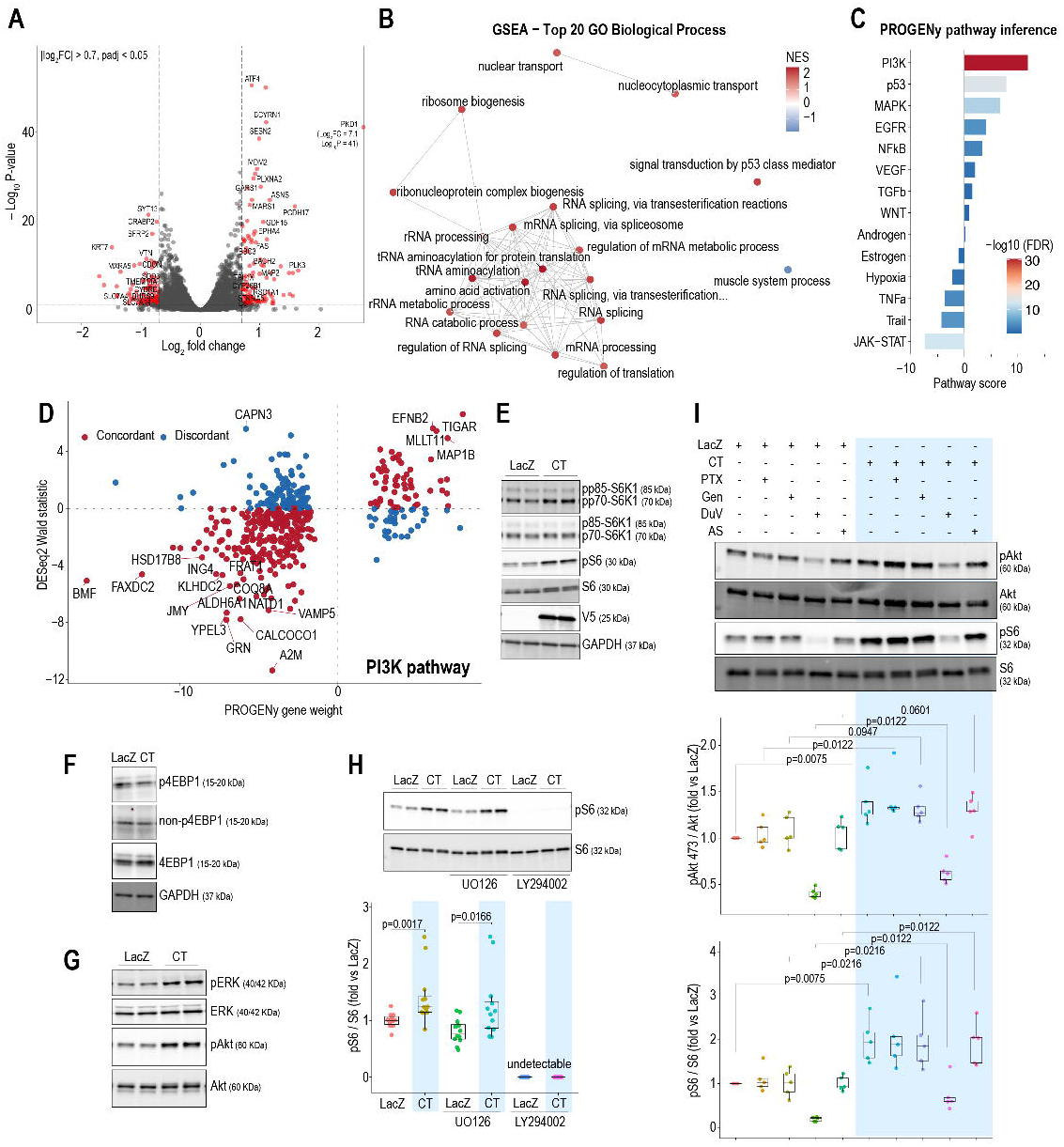
Effect of PC1-CT overexpression on gene expression and protein synthesis-related pathways in hiPSC-CMs. **A**. Volcano plot from bulk RNA-seq showing global gene expression changes in PC1-CT overexpressing hiPSC-CMs compared to LacZ control. A total of 105 genes were significantly downregulated, and 102 were upregulated (red circles indicate genes with |log_2_FC| > 0.7 and FDR < 0.05). **B**. Gene Set Enrichment Analysis (GSEA) using the GO: BP gene set. RNA-seq data reveals significant enrichment of gene sets related to protein synthesis, ribosome biogenesis, RNA processing, and nuclear export. **C**. Pathway inference using decoupleR and PROGENy using DESeq2 Wald statistic showing upregulated PI3K pathway. **D**. PROGENy gene weights for the inferred upregulated PI3K pathway. **E**. Representative Western blots showing increases in p70-S6K1 and S6 phosphorylation after PC1-CT overexpression compared to LacZ control. V5 was used to confirm PC1-CT overexpression. **F**. Representative Western blots show no changes in 4EBP1 phosphorylation among groups. **G**. PC1-CT increases both ERK and Akt phosphorylation. **H**. Inhibition of PI3K/Akt using LY294002 blunted p-S6, whereas the MEK inhibitor U0126 had no effect (N=12 biological replicates from 6 independent differentiation batches). **I**. Effect of pertussis toxin (PTX), genistein, and PI3Kγ inhibitors on p-S6 and p-Akt (N=5 biological replicates from 5 independent differentiation batches). GAPDH is shown as a loading control for panels **E** and **F**. A Mann-Whitney test was used to compare groups, as indicated in Figures **H-I**.

### PC1-CT activates PI3K-Akt-mTOR-S6K1-S6 signaling

mTOR is a key regulator of cardiomyocyte protein synthesis [17]. Because RNA-seq indicated enrichment of protein synthesis-related pathways, we examined mTOR signaling by assessing phosphorylation of S6K1 and S6. PC1-CT overexpression increased phosphorylation of p70-S6K1 and S6 relative to LacZ (**Fig. 2E**). In contrast, 4EBP1 phosphorylation status was unchanged (**Fig. 2F**), suggesting that PC1-CT may preferentially activate the S6K1-S6 branch of mTOR signaling. Because mTOR can be activated through both PI3K-Akt and MEK-ERK pathways [17], we next assessed Akt and ERK phosphorylation. Both pathways were activated in PC1-CT relative to LacZ (**Fig. 2G**). To determine the contribution of these pathways to S6 activation, we treated cells overnight with the pan-PI3K inhibitor LY294002 (50 μM) or the MEK inhibitor U0126 (10 μM). LY294002 abolished the increase in pS6, whereas U0126 had no effect (**Fig. 2H**), indicating that PC1-CT-induced S6 activation is predominantly PI3K-dependent.

Because PC1-CT has been reported to engage G_i/o_-dependent signaling [4], we first tested whether its effects on Akt/S6 required G_i/o_ activity using pertussis toxin (1 μg/mL, 6 h). To assess potential alternative tyrosine kinase-dependent mechanisms, we also applied the broad tyrosine kinase inhibitor genistein (100 μM, 6 h). Finally, because PI3Kγ is classically linked to G_i/o_-coupled signaling, we evaluated the effects of the small-molecule inhibitors duvelisib and AS-605240 (1 μM each, 6 h). Pertussis toxin, genistein, and AS-605240 did not alter PC1-CT-induced pAkt or pS6 levels, whereas duvelisib modestly reduced basal signaling (**Fig. 2I**). Together, these data suggest that PC1-CT-induced Akt/S6 activation is largely independent of G_i/o_ signaling and PI3Kγ.

## DISCUSSION

Previous work in rodent cardiomyocytes has implicated PC1-CT in cardiomyocyte hypertrophy [3, 4]; however, its localization and function in human cardiomyocytes remain undefined. We identify a striated, largely sarcomeric pool of PC1 C-terminal signal in cardiomyocytes and show that PC1-CT overexpression recapitulates this localization, activates PI3K-Akt-mTOR-S6K1-S6 signaling, and increases ribosome biogenesis and protein synthesis-related gene expression.

PC1 association with the actomyosin complex and focal adhesion complexes has been described [18, 19], but, to our knowledge, its localization in cells with organized sarcomeres has not. The endogenous PC1 C-terminus in hiPSC-CMs displayed a regular striated pattern with minimal overlap with the sarco-endoplasmic reticulum marker KDEL. It remains unknown whether this signal represents a cleavage fragment containing only the PC1-CT. Notably, adenoviral overexpression of PC1-CT in hiPSC-CMs and adult mouse ventricular cardiomyocytes generated a similar striated distribution, indicating that PC1-CT (aa 4102–4303) alone is sufficient to confer this localization pattern. In hiPSC-CMs, localization occurred between the α-actinin bands. By contrast, in adult mouse ventricular cardiomyocytes, overexpressed PC1-CT showed greater overlap with the sarcomeric Z-disc proteins α-actinin and desmin. This difference suggests a maturation-dependent localization pattern. Additional studies are required to define the precise sub-sarcomeric localization and identify the relevant interacting partners.

PC1-CT overexpression induced gene expression programs enriched for ribosome biogenesis, RNA processing, nuclear export, and protein synthesis-related pathways, while classical hypertrophic markers remained unchanged. These transcriptomic changes coincided with increased phosphorylation of Akt, S6K1, and S6, whereas 4EBP1 phosphorylation remained unchanged, indicating preferential engagement of the S6K1-S6 branch rather than global mTOR activation. Additionally, LY294002, but not U0126, suppressed S6 phosphorylation, indicating that PI3K lies upstream of S6 activation. Together, these findings support a model in which PC1-CT activates a PI3K-Akt-mTOR-S6K1-S6 pathway that drives biosynthetic remodeling without evidence of a canonical fetal-gene hypertrophic response over the time frame examined.

Because LY294002 can inhibit mTOR in biochemical assays [20], we interpret this pharmacological result cautiously; however, LY294002 does not directly inhibit mTORC2-dependent Akt phosphorylation [21]. Additional evidence links PC1 to mTOR signaling. Tuberous sclerosis complex 2 (TSC2) is a key repressor of mTOR and lies adjacent to *PKD1* on chromosome 16p13.3. Contiguous germline *TSC2* and *PKD1* deletions cause severe infantile-onset polycystic kidney disease [22, 23]. PC1-CT has also been reported to bind TSC2 [24], and PC1 has been linked to the regulation of Akt-mTOR [25] and ERK-mTOR signaling [26]. While activation of mTOR has been observed in renal cysts from ADPKD patients and mouse models [24, 27], *Pkd1*^-/-^ mouse embryonic fibroblasts did not have evidence of constitutive mTOR activation [28], suggesting that downstream consequences of PC1-CT signaling are likely cell-type- and stimulus-dependent.

These findings extend prior work showing that a synthetic membrane-anchored PC1-CT construct activates Akt via a pertussis toxin-sensitive, G_i/o_-dependent pathway in cardiomyocytes [4]. Our data show that non-anchored PC1-CT overexpression mirrors endogenous localization and can activate Akt despite insensitivity to pertussis toxin and PI3Kγ inhibitors. This difference may reflect distinct signaling outputs of membrane-anchored and non-membrane-anchored PC1-CT constructs, potentially separating regulation of LTCC trafficking from biosynthetic signaling.

This study has several limitations. Most importantly, the mechanistic experiments rely on PC1-CT overexpression, which may not fully recapitulate endogenous cleavage, trafficking, abundance, or stoichiometry. We did not directly compare membrane-anchored and non-membrane-anchored PC1-CT constructs in the same system, identify the upstream PI3K isoform(s) or proximal interacting proteins responsible for signaling, or resolve the precise sub-sarcomeric localization of PC1-CT. Future studies should define endogenous PC1 cleavage and trafficking to the sarcomere, map PC1-CT binding partners, and determine whether physiological or pathological mechanical stress modulates this pathway. These efforts will be essential to fully elucidate the role of PC1 in human cardiomyocytes.

## ACKNOWLEDGMENTS

We thank Dr. Peter C. Harris and the Mayo PKD Center (Division of Nephrology and Hypertension, Mayo Clinic) for sharing the mCherry-PC1 plasmid; Dr. Joseph A. Hill (Division of Cardiology, UT Southwestern) for the PC1-CT adenovirus; and the Advanced Cellular and Tissue Microscopy Core and Research Pathology Core at Houston Methodist.

## DECLARATION OF AI-ASSISTED TECHNOLOGIES IN THE WRITING PROCESS

During the preparation of this work, the author(s) used OpenAI models to improve readability and grammar. After using this tool/service, the author(s) reviewed and edited the content as needed and take full responsibility for the content of the publication.

## SUPPLEMENTAL TABLE LEGENDS

**Supplemental Table 1: Differentially expressed genes in hiPSC-CMs infected with LacZ and PC1-CT adenovirus.** Comparison between groups was performed using DESeq2 (Wald test p-value and Benjamini-Hochberg adjusted p-value).

